# Predictive evolution of metabolic phenotypes using model-designed selection niches

**DOI:** 10.1101/2021.05.14.443989

**Authors:** Paula Jouhten, Dimitrios Konstantinidis, Filipa Pereira, Sergej Andrejev, Kristina Grkovska, Payam Ghiachi, Gemma Beltran, Eivind Almaas, Albert Mas, Jonas Warringer, Ramon Gonzalez, Pilar Morales, Kiran R. Patil

## Abstract

Traits lacking fitness benefit cannot be directly selected for under Darwinian evolution. Thus, features such as metabolite secretion are currently inaccessible to adaptive laboratory evolution. Here, we utilize environment-dependency of trait correlations to enable Darwinian selection of fitness-neutral or costly traits. We use metabolic models to design selection niches and to identify surrogate traits that are genetically correlated with cell fitness in the selection niche but coupled to the desired trait in the target niche. Adaptive evolution in the selection niche and subsequent return to the target niche is thereby predicted to enhance the desired trait. We experimentally validate the theory by evolving *Saccharomyces cerevisiae* for increased secretion of aroma compounds in wine fermentation. Genomic, transcriptomic, and proteomic changes in the evolved strains confirmed the predicted flux re-routing to aroma biosynthesis. The use of model-designed selection niches facilitates the predictive evolution of fitness-costly traits for ecological and biotechnological applications.

## Introduction

Adaptive evolution can give rise to and optimize complex phenotypes as evident from the current and historical biodiversity (*1–4*). Evolutionary optimization can also be observed in molecular detail in the laboratory, as illustrated by, for example, the *Escherichia coli* long-term evolution experiment (*5*). While the evolution can involve numerous alternative paths at the level of the genotype, Darwinian selection constrains the phenotypic end points (fitness) (*6*). The phenotypic evolution can therefore be more precisely controlled through environmental selection pressure. This allows improving cells without the explicit knowledge of the underlying genotype-phenotype relations using adaptive laboratory evolution. Adaptive laboratory evolution has been used for improving microbial strains, including alleviation of fastidious vitamin requirements (*7*), and enhancement of photosynthetic capabilities of engineered microorganisms (*8, 9*).

While useful in optimizing complex traits and operationally simple, adaptive laboratory evolution is inherently limited to the traits genetically linked to the cell fitness. Improving fitness-neutral or costly trait requires artificial, non-Darwinian, selection through screening of large numbers of variants. This is a considerable combinatorial challenge for complex multigenic traits, and, thus, application of artificial selection has yet been limited to single proteins or pathways with photometric readouts (*10–14*). Adaptive selection of a complex trait requires identification of an environmental condition where the trait becomes growth-linked (*15*), for example, selection for increased antioxidant production under oxidative conditions (*16*). However, generalization of such anecdotal and intuition-guided methods is difficult and calls for predictive models of trait dependences.

Here, we ask whether first-principle models could enable use of Darwinian selection for enhancing traits that carry no fitness benefit. We base our strategy on genome-scale metabolic models, which allow predicting metabolic fluxes consistent both with the mass balance constraints and the fitness objectives of the cells (e.g. optimal growth) (*17, 18*). In the context of laboratory evolution, genome-scale metabolic models have been shown to predict fitness improvement and the associated metabolic flux changes (*19–22*). We use these models to predict environment-dependence of the correlations between metabolic traits, and that between metabolic traits and the cell fitness. This allowed us to predict selection environments, distinct from the target niche wherein the trait manifestation is required, for adaptive laboratory evolution. We put the predicted selection niches to experimental test through the improvement of wine aroma production in yeast.

## RESULTS

### Selection niche

Consider a target niche wherein the manifestation of a metabolic trait of interest is desired, e.g. wine must. We propose that an improvement of the desired trait, e.g. aroma production, in the target niche can be achieved through adaptive evolution in a distinct selection niche followed by the return to the target niche. To devise the selection niche, we take advantage of the observation that the covariance between any two traits is dependent on the growth environment.

The selection acting on a phenotypic trait is the covariance between the trait and the relative fitness, as described by Robertson-Price identity (*23–26*) (Equation 1).

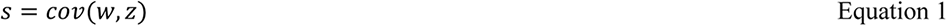

where *s* is the selection differential, *w* fitness, and *z* the trait of interest.

When there is genetic covariance between the trait and relative fitness, evolutionary response to selection can occur (Equation 2, the secondary theorem of selection).

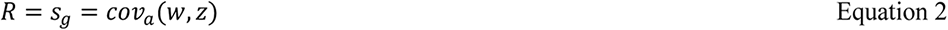

where *R* is the response to selection, *s_g_* is the genetic selection differential, and *cov_a_(w,z)* is the additive genetic covariance. Equation 2 generalizes to a multivariate form for multiple traits (*27*).

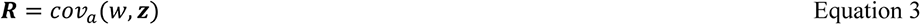

A metabolic trait can be modelled as a set of metabolic fluxes. Metabolic trait interdependencies (i.e. flux couplings) under a given nutrient environment can then be predicted using genome-scale metabolic models (*28*). Two metabolic reactions are coupled if a non-zero flux through one reaction implies a non-zero flux through the other. For an organism adapted to a particular environment, flux coupling leads to genetic dependences between the corresponding enzyme-coding genes (*29*). We use the coupling between metabolic traits and the specific growth rate (proxy for mean fitness) to predict relative responses to selection analogously to the secondary theorem of selection (Equation 4).

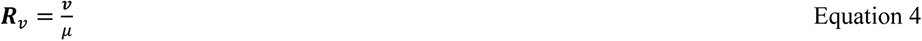

where ***R****_v_* is the relative responses of single-flux metabolic traits to selection, *v* the metabolic fluxes, and *µ* the specific growth rate.

We use the selection response relation (Equation 3) as a basis to search for a suitable selection niche – a defined chemical environment in which the adaptive evolution is to take place. Ideally, the selection niche would be chosen such that there is a direct selection for the desired trait through coupling with cell growth. This, however, will only rarely be possible as most desired traits, such as metabolite secretion, are a trade-off with cell growth due to competition for metabolic precursors and co-factors (*30, 31*) (Fig. 1a). We therefore aim at growth coupling of a surrogate trait, which we define as a set of fluxes that are coupled to cell growth in the selection niche, and with the desired trait in the target niche (Fig. 1b). It is not necessary for the surrogate trait to be coupled with the desired trait in the selection niche, nor it is likely, due to the trade-off against the growth. Thus, the surrogate trait is necessarily a proper subset of fluxes that must increase or decrease for the desired trait enhancement in the target niche. Due to the dependence of genetic correlations between traits on the growth environment (Equation 3), the surrogate trait and the selection niche are inherently linked to each other and need to be identified simultaneously.

**Figure 1.**
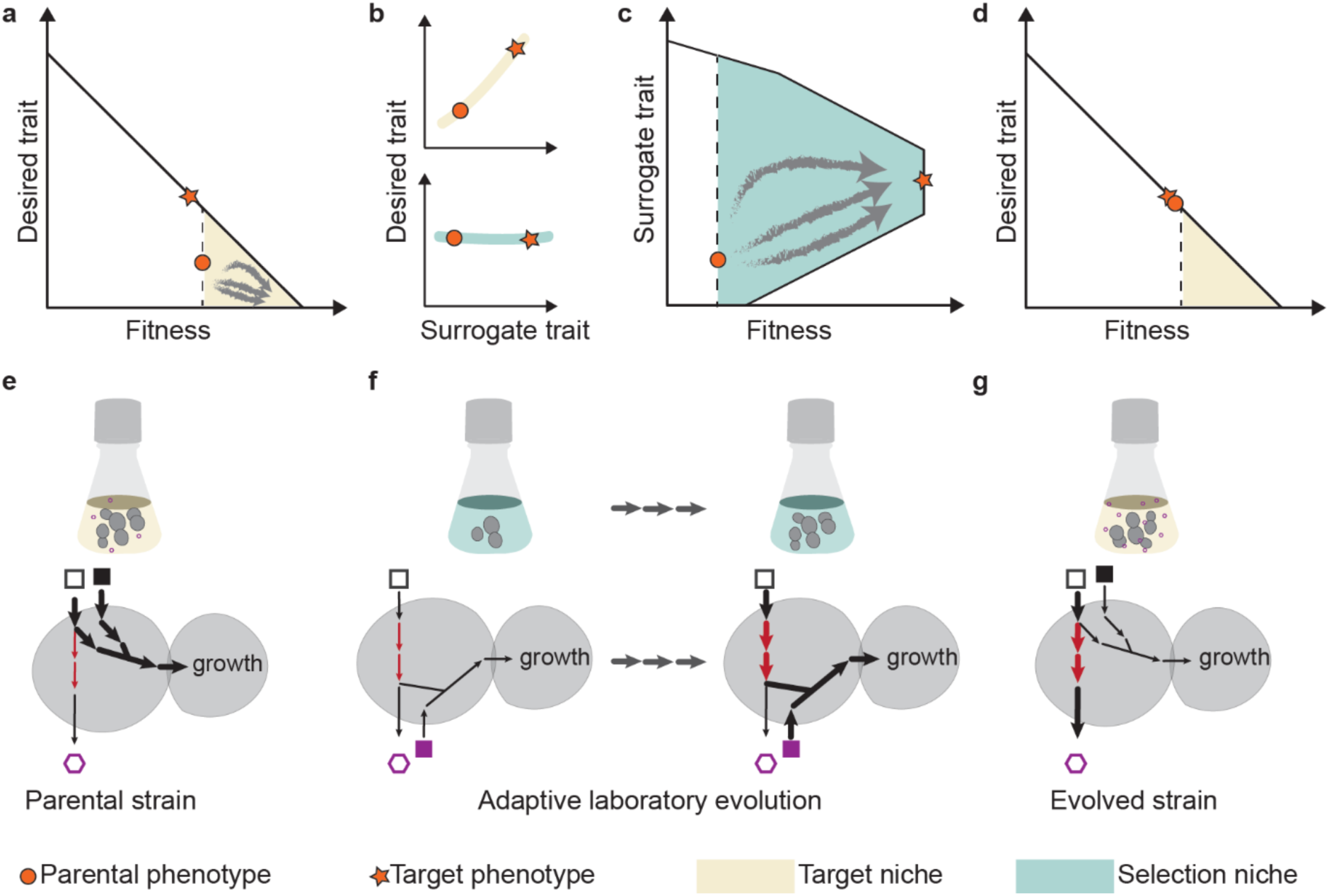
Selection niches and surrogate traits for Darwinian selection in absence of fitness advantage. Current phenotype is represented with an orange circle whereas the orange star represents a desired phenotype. a) In the target niche (yellow), Darwinian process (grey arrows) selects for fitter phenotypes with diminished desired trait(s). b) The surrogate trait is chosen such that it is coupled to the desired trait in the target niche, but not in the selection niche (green). c) The surrogate trait is coupled to fitness in the selection niche and can therefore be improved through Darwinian selection. d) Evolved cells with a strengthened surrogate trait manifest an improved desired trait in the target niche. e) A simplified schematic example of how the selection niche can be used for evolving desired traits. The desired trait in the example is the production of the open hexagon. The squares are nutrients available differing between the target and selection niches. The arrows represent metabolic fluxes, the thicker the arrow the higher the flux. The desired trait does not share metabolic fluxes (i.e. genotypic features) with growth (i.e. proxy for mean fitness) in the target niche, but the surrogate trait (red arrows) is a subset of the flux basis of the desired trait. In the selection niche, the surrogate trait is coupled to fitness through shared intracellular fluxes. Thus, the surrogate trait can be improved through adaptive evolution in the selection niche. The improved surrogate trait supports the desired trait in the target niche.

In the target niche, a desired trait that does not pose a fitness advantage will not be under Darwinian selection (Figure 1a). But the selection niche is designed such that it couples the surrogate trait to mean fitness (Figure 1c), allowing positive selection on *de novo* mutations enhancing the surrogate trait. Following environment switch to the target niche, in which the surrogate trait is coupled with the desired trait, the latter is enhanced (Figure 1d and 1e). To visualize this process, consider a simple metabolic network (Figure 1e-g). While the parental strain is well adapted to channel the nutrients to cell growth (Figure 1e), in the selection niche different metabolic fluxes are coupled with growth (Figure 1f). When the selection niche is designed to couple the surrogate trait to growth, the mutations enriched in the selection niche improve the desired trait in target niche (Figure 1g). While the desired trait may be negatively selected in the target niche, this is not an obstacle for use in applications. A typical microbiological process involves only a few generations (below ten), and is thus unlikely to diminish the desired trait. The necessary condition will be to maintain the cell stock in a separate environment (in this case the selection niche), which is a common practice in microbiological applications.

### Predicting selection niche

To predict selection niches satisfying the conditions laid out above, we devised an algorithm, termed EvolveX, based on genome-scale metabolic models. In brief, the algorithm simultaneously identifies a surrogate trait and evaluates the suitability of a niche – a set of available nutrients – as a selection niche for adaptively evolving the surrogate trait.

The EvolveX algorithm involves four principle steps. *Step 1*: The flux basis of the desired trait is determined as fluxes that must change the desired trait to become enhanced in the target niche. *Step 2*: A response to selection of the desired trait’s flux basis is predicted in the selection niche. The subset of the flux basis with predicted non-zero responses to selection forms the surrogate trait. As the covariances of traits may change through evolution (*32–34*), we define the flux basis of the desired trait (*Step 1*) in the ancestral state but predict the response to selection in an evolved state that is expected to be approached during experimental evolution (*19, 22, 35*). *Step 3*: A minimum size (cardinality) of the flux basis subset having stronger desired response to selection in the selection niche than in the target niche is estimated. *Step 4*: A suitability score of a selection niche is calculated by combining: (a) results of *Step 1*, indicating the strength of response to selection; (b) results of *Step 2*, indicating the coverage of flux basis with desired selection; and (c) the number of chemical components in the selection niche. The last criterion is included to discount for the uncertainty in the knowledge concerning the organism’s nutritional preferences. The details of EvolveX implementation, including accounting for alternative steady-state flux distributions, and normalization to enable comparison across different growth media likely eliciting different growth rates, are provided in Methods.

### Model-predicted selection niches increase aroma production

To experimentally test the concept of the selection niche in adaptive evolution, we used EvolveX to improve secretion of aroma compounds by *S. cerevisiae* in wine must, a niche that is characterized by high sugar content and relatively less assimilable nitrogen including amino acids. As aroma synthesis diverts carbon and nitrogen away from the production of daughter cells, it cannot be selected in adaptive evolution in the target niche (wine must). Furthermore, while the metabolic pathways that synthesize aroma compounds are known, their regulation is poorly understood, preventing facile engineering of aroma secretion (*36*). We targeted two main groups of aroma compounds for selection: phenylethyl alcohol and the derived acetate ester, phenylethylacetate, which have a rose and honey scent and raspberry-like flavor; and, branched-chain amino acid-derived higher alcohols (i.e. 2-methyl-1-butanol and 3-methyl-1-butanol) and their acetate esters (i.e. 2-methylbutylacetate and isoamyl acetate), which have a banana and pear scent and fruity flavor.

To identify a suitable selection niche and surrogate trait, we assessed all 1540 combinations of up to three carbon and nitrogen sources, out of 22 common constituents of yeast growth media. All potential niches were ranked by using the EvolveX score (Step 3) (supplementary information, Table S1). Following literature-based assessment of substrate utilization preferences of *S. cerevisiae*, two of the high-scoring niches (in the top 20 out of the 1171 niches that supported growth) were selected for experimental evolution. A selection niche containing glycerol, phenylalanine, and threonine as the sole carbon and nitrogen sources was chosen for phenylethyl alcohol and phenylethylacetate production, creating a surrogate trait of seven fluxes out of the flux basis of 20 (supplementary information, Table S2). This growth medium is referred to as the glycerol niche (Figure 2a). For increasing the production of branched-chain amino acid-derived aromas, a selection niche containing ethanol, arginine, and glycine was chosen, creating a surrogate trait of eleven fluxes out of the flux basis of 44 (supplementary information, Table S2). This growth medium is referred to as the ethanol niche (Figure 2a).

**Figure 2.**
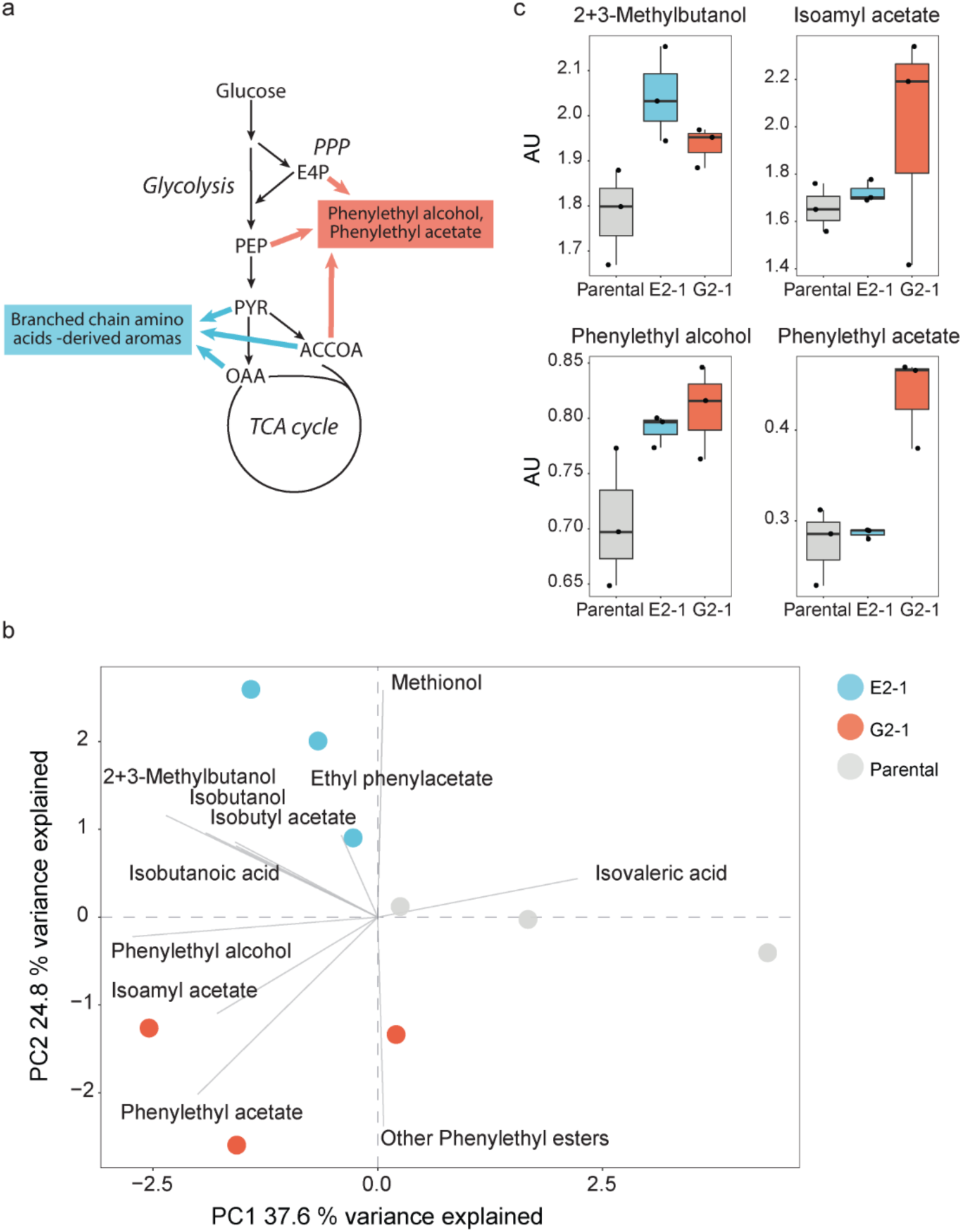
Adaptive laboratory evolution of *S. cerevisiae* in model-designed selection niches led to predicted changes in aroma compound production in wine must fermentations. a) Origin of aroma compounds in the yeast central metabolism: branched-chain amino acid derived compounds (esp. 2-methyl-1-butanol, 3-methyl-1-butanol, isoamyl acetate and 2-methylbutylacetate), and aromatic amino acid derived compounds (esp. phenylethyl alcohol and phenylethyl acetate). Acetate esters of higher alcohols share an acetyl-CoA (ACCOA) precursor. b) Principal components analysis of aromatic and branched amino acids-derived aroma compound profiles of natural wine must fermentations, with the parental (grey) and evolved strains. Strain from the ethanol selection niche (ethanol, arginine, glycine in light blue, that from the glycerol selection niche (glycerol, phenylalanine, threonine) in orange. First principle component mainly captures the variation between the parental strains and the evolved strains while the second principle component captures the variation between the isolates from the two different selection niches. c) Selected aroma compound abundances in wine must fermentations of evolved strains (the scale is in arbitrary units, AU). E2-clone 1 (light blue) was selected in the ethanol niche, and G2-clone 1 (orange) was evolved in the glycerol niche. 2+3-methylbutanol (a combined pool of 2-methyl-1-butanol and 3-methyl-1-butanol) and isoamyl acetate (acetate ester of 3-methyl-1-butanol) were the desired target aromas of the ethanol niche, deriving from branched-chain amino acids. Phenylethyl alcohol and its acetate ester, phenylethyl acetate, were the desired target aromas of the glycerol niche.

A diploid *S. cerevisiae* wine strain was grown in the two selection niches. Three replicate populations were independently evolved asexually for ∼150 generations. The parental strain and a single colony isolate (out of nine) from all evolved populations were then characterized in industrially relevant wine must fermentations. The fermentation performance of the selected strains was monitored through culture weight loss and the HPLC profiles of sugars and the main fermentation byproducts (i.e. ethanol, glycerol, acetate) of the cultures (supplementary information, Table S3). The evolved isolates maintained their fermentation performance in wine must indicating their suitability for use in wine fermentations. The volatile flavor compounds in the wine must fermentations were quantified using GC-MS analysis. Principal component analysis of the quantified volatile compounds of interest, those deriving from aromatic (four compounds or indistinguishable compound sets) and branched chain amino acids (seven compounds or indistinguishable compound sets), showed a clear divergence of the evolved strains from the parental aroma profile (Figure 2b). Further, the isolates selected in the glycerol niche were distinct from those selected in ethanol niche. The volatile compounds separating the three groups were in accord with the model predictions. The first principle component (PC1, 38% of the total variance) distinguished the parental strains from the evolved strains, the separation being driven by the surrogate traits that overlapped between the two selection niches – fluxes through transketolase 1 and ribulose 5-phosphate 3-epimerase. In accord with the model predictions, the isolates selected in the ethanol and glycerol selection niches were separated by the target aroma compounds except isoamyl acetate (PC2, 25% of the total variance, Figure 2b-c). Phenylethylacetate was specifically increased in fermentations with the isolates selected in the glycerol niche (Figure 2c). Similarly, the combined pool of branched-chain amino acid-derived aroma compounds 2-methyl-1-butanol and 3-methyl-1-butanol was increased only for the isolate selected in the ethanol niche.

### Evolved strains exhibit molecular changes in line with the model predictions

To understand the genetic basis of the selected phenotypes, we sequenced the whole genomes of the evolved populations. In addition, we sequenced the isolates from the evolved populations at an earlier (∼100 generations) and at a later (∼214 and 165 generations in ethanol niche and in glycerol niche, respectively) stage of the adaptive laboratory evolution process. The clones and the populations selected in the glycerol niche experienced low frequencies of single nucleotide variants (SNVs) and no genes showed recurrent SNVs. However, copy number variants (CNVs) were prevalent with a majority of the genome having been triplicated in some populations and clones (Figure 3a; supplementary information, Table S4).

**Figure 3.**
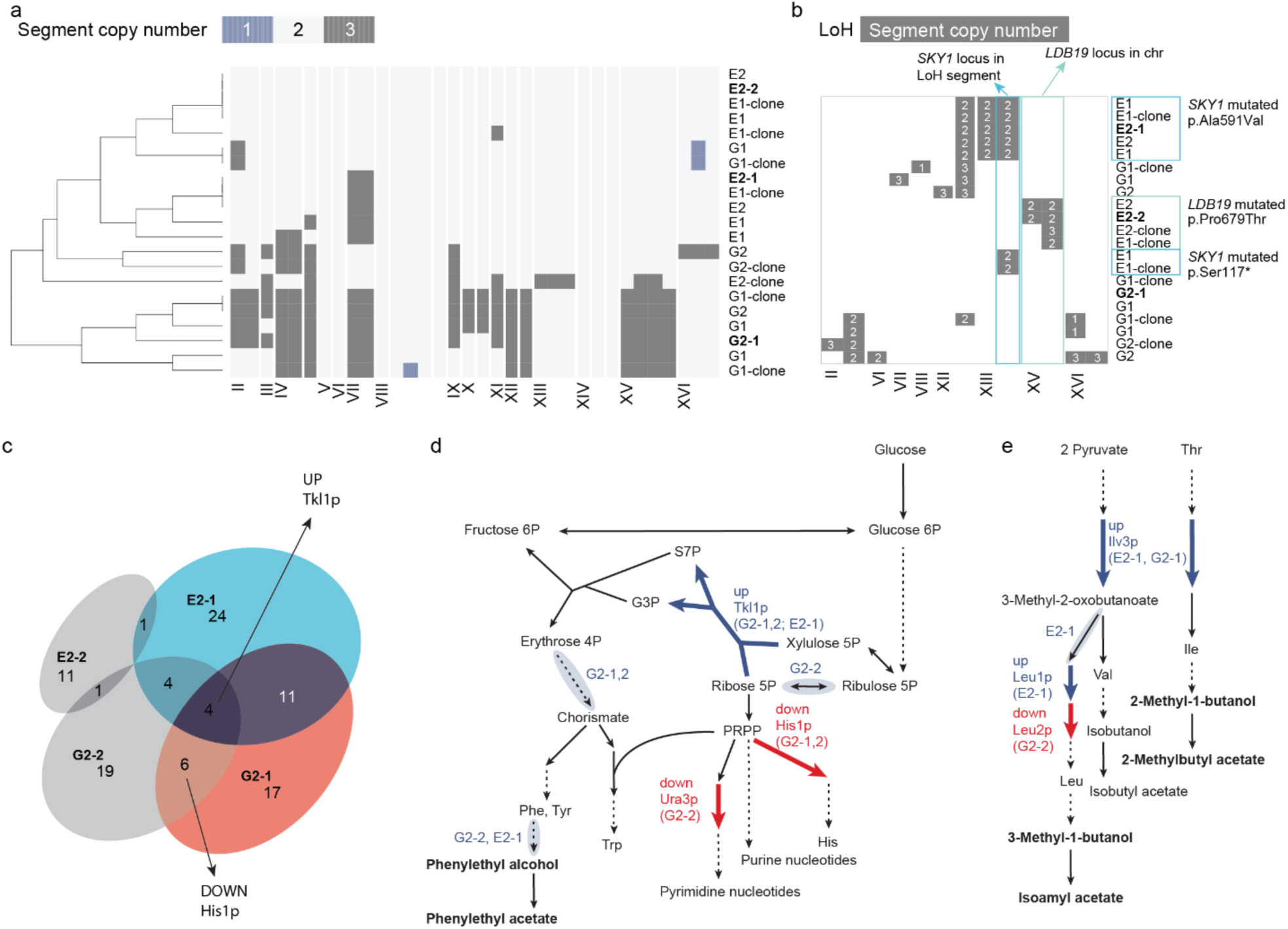
Evolved yeast strains show molecular changes in line with the model predictions. a) Evolved populations and clones from the glycerol niche (indicated with G) exhibited large CNVs. Genome segment copy numbers are shown for evolved populations and clones (from glycerol and ethanol niche indicated with G and E, respectively) along the chromosomes. Vertical lines indicate ends of contigs. The clones for which we determined protein and transcript alterations are indicated in bold. b) Loss-of-heterozygosity was associated with SNVs in evolved populations, and clones from the ethanol niche. c) Proteomics analysis of the evolved clones grown in wine must (the target niche) revealed both shared and niche-specific changes in comparison to the parental strain (three biological replicates, multiple testing corrected *p*-value < 0.01, -1 ≥ log2 fc ≥ 1). Clones for which we measured the aroma production are highlighted with blue and orange. Clones from the glycerol niche specifically featured higher abundance of Tkl1p and lower abundance of His1p. d) Evolved clones grown in wine must featured changes in protein and transcript abundances (three biological replicates, fdr < 0.05, -1 > log2 fc > 1) in the pathways leading to the target aroma compounds phenylethyl alcohol and phenylethyl acetate. The changes consistent with the predictions in specific clones are indicated with colored arrows (protein-level) and clouds around the arrows (transcript-level). e) Proteomic and transcriptomic changes in evolved cells, marked as in d), for pathways leading to the branched chain amino acids derived target aroma compounds.

In contrast, in populations and clones selected in the ethanol niche the chromosome copy number aberrations were limited to chromosome VII (triplicated), and recurrent SNVs were found in *SKY1*, encoding a protein serine/threonine kinase involved in the regulation of polyamine transport (missense p.Ala591Val, frameshift p.Leu64fs, stop gain p.Ser117*). We also observed a loss-of-heterozygosity (LoH) segment in the contig containing the *SKY1* locus (supplementary information, Table S4). *SKY1* deletion gives yeast tolerance to high spermine concentrations (*37*), a degradation product of arginine, one of the three components in the ethanol niche. In cases where *SKY1* mutations were not detected, we either found paired missense mutations in genes encoding the ubiquitin ligase Rsp5 (p.Arg355Gly) and its target-guide and adapter Ldb19 (p.Pro679Thr), which drive the endocytosis of plasma membrane-localized amino-acid transporters, and a LoH in the contig containing the *LDB19* locus (supplementary information, Table S4). Thus, in the ethanol niche, mutations in genes involved in arginine utilization were enriched during adaptive laboratory evolution, consistent with the selection regime. However, the recurrent copy number variants enriched in the glycerol niche affected multiple genes rendering it challenging to elucidate associations with either the particular selection or the improved aroma generation trait.

To better understand the complex multigenic changes associated with the improved aroma production phenotypes, we investigated the underlying transcript and protein abundance states. We analyzed these in wine must (our target niche), as well as in the selection niches. The overlap of differentially expressed transcripts and proteins was limited (max 6 % of all differentially abundant proteins and RNAs in a strain) suggesting post-transcriptional regulation plays a substantial role in both the improved aroma generation in wine must and to the improved fitness in the selection niches. The fitness improvement in the glycerol niche was associated with a differential abundance of 40 (evolved strain G2-2) and 38 (evolved strain G2-1) metabolic enzymes (as per yeast genome-scale metabolic model v. 7.6; Aung, Henry and Walker, 2013). The proteins with increased abundance were enriched in respiratory processes: oxidative phosphorylation for G2-1, (*p*-value 2.74e-05) and mitochondrial electron transport for G2-2 (*p*-value 0.0424). This is in accord with the strong selection pressure on these processes, as predicted by EvolveX (supplementary information, Table S5). Additionally, three glycolytic enzymes (Cdc19, Pdc6, and Tdh1) become less abundant in G2-1, suggesting increased respiratory activity relative to glycolysis.

The fitness improvement in the ethanol niche was also associated with only a handful of enzyme abundance changes: 13 in evolved strain E2-2 and 10 in evolved strain E2-1 (supplementary information, Table S5). Consistent with arginine being the sole nitrogen source in this selection niche, these changes involved arginine metabolism, including decreased abundance of arginine biosynthetic pathway enzymes (Arg1 and Arg8 in E2-1, and Arg5,7 in E2-2). Strain E2-2 further had decreased abundance of proline oxidase Put1, involved in the utilization of one of the four nitrogen atoms in arginine. Several transporters had higher abundance in E2-2: arginine permease (Can1), monocarboxylate transporter (Jen1), methionine permease (Mup1), and hexose transporter (Hxt6). The endocytosis of all of these transporters is mediated by Rsp5-Ldb19 (*39–41*), which is mutated in the E2-2. Thus, in both selection niches, the changes in protein expression were limited and included pathways predicted by EvolveX – respiration in the case of the glycerol niche and arginine metabolism in the case of the ethanol niche.

In the target niche, wine must, the desired traits of improved aroma generation were associated with few changes in protein abundances (11–24 proteins, Figure 3c). Changes in metabolic enzymes could be attributed to increased target aroma precursor supply or to decreased competition for the precursors. The precursor supply was enhanced in all evolved strains through increases in the level of transketolase (Tkl1), which was included in the both surrogate traits (i.e. for increasing the aromatic and branched chain amino acid derived aromas). The clones from the glycerol niche specifically showed decreased levels of ATP phosphoribosyltransferase His1, which competes with Tkl1 for the precursor ribose 5-phosphate (Figure 3d). Orotidine-5’-phosphate decarboxylase (Ura3) involved in purine nucleotide synthesis, was found in decreased abundance in one of the clones from the glycerol niche, consistent with aroma synthesis competing for nucleotide precursors. Gene expression changes influencing precursor supply were observed in the clones from the glycerol niche, and specifically genes involving chorismate synthesis (*ARO1*, *ARO3*, and *RKI1* being part of the trait flux basis), in the gene for aromatic amino transferase (*ARO8*), and in the *PDC1*, *PDC5* genes, which encode phenylpyruvate decarboxylating enzymes involved in the Ehrlich pathway of phenylethyl acetate formation (*42*) and belong to the surrogate trait.

While the clones from the ethanol niche and the glycerol niche both showed increased abundance of Tkl1, the ethanol niche clones did not show reduced levels of His1 or Ura3 (Figure 3c). Instead, one of the ethanol niche clones showed increased levels of dihydroxyacid dehydratase (Ilv3) and isopropylmalate isomerase (Leu1), involved in branched-chain amino acid biosynthesis, the origin of the target aroma compounds (Figure 3e). Both Ilv3 and Leu1 belonged to the corresponding trait flux basis, as does beta-isopropylmalate dehydrogenase (Leu2), which catalyzes the reaction following that catalyzed by Leu1 in the leucine biosynthesis pathway. However, the flux basis of enhanced phenylethyl acetate generation was coupled with Leu2 down-regulation, and one of the clones from glycerol niche showed the predicted decrease in Leu2 abundance (Figure 3e, supplementary information, Table S5). Thus, we were able to directly associate altered protein abundances in the clones selected in the glycerol and ethanol niche with the corresponding desired aroma synthesis pathways and the competing metabolic pathways.

## Discussion

In our approach, we are able to select for what would otherwise be fitness-costly or fitness-neutral traits by placing the cells in an appropriately chosen selection niche. Our theory and experimental results show that this selection niche can be predicted using first-principle modelling. We used such model-designed niches to improve two aroma production traits in wine yeasts. Previously, adaptive evolution of fitness-beneficial traits in one niche has been shown to facilitate exaptation, i.e. the predisposition to fitness improvement in another niche (*22*). In contrast, we propose and show that, in an appropriately chosen selection niche, a trait without a fitness benefit in the target niche can adaptively evolve.

Using surrogate traits and the selection niche maintains the key advantage of adaptive laboratory evolution, in circumventing the need to know the genetic and regulatory basis of the traits of interest beyond the basic metabolic network structure. Indeed, analysis of the improved aroma generation traits in our case study revealed complex genotype-phenotype relationships. Improving these traits using rational strain improvement would currently be challenging (Hassing *et al.*, 2019). Our approach of combining surrogate traits with the selection niche makes such traits amenable for adaptive evolution. As genome-scale metabolic models are becoming easier to reconstruct (*44–47*), our approach can be applied to any organism amenable for experimental evolution.

The increased phenylethyl alcohol and phenylethyl acetate generation we observed occurred in a selection niche containing the direct aroma precursor phenylalanine. In contrast, no target aroma precursor was included in the selection niche for the branched-chain amino acid-derived aroma compounds. This demonstrates the utility of metabolic modelling in identifying non-intuitive selection niches. While we designed the selection niches in this study using carbon and nitrogen sources, enzyme inhibitors and substrate analogs can also be used to expand the search space for the selection niches.

The theory of the surrogate traits and the selection niches is generalizable to non-metabolic traits as long as the dependency between the fitness and other traits can be quantitatively modelled. This could be done by using signaling networks or genetic interaction networks, for example. Our theory provides means for understanding complex adaptive processes through systematizing the joint environment-genetic dependences between fitness and other traits, and also enables the manipulation of complex traits for biotechnological applications.

## Author contributions

Conceptualization PJ, EA, KRP; Methodology PJ, KRP; Software PJ; Investigation PJ, DK, PM, FP, KG, PC; Formal Analysis PJ, DK, PM, FP; Resources RG, SA, GB, AM; Visualization PJ and KRP; Writing – Original Draft PJ and KRP; Supervision KRP, RG, JW, AM, EA.

## Acknowledgements

This work was sponsored by the ERASysAPP project WINESYS (the German Ministry of Education and Research grant no. 031A605; the Research Council of Norway grant no. 245160) and by the Ministry of Science, Innovation and Universities, Spain (Project CoolWine, PCI2018-092962), under the call ERA-NET ERA COBIOTECH. PJ acknowledges funding from the Academy of Finland, decision numbers 310514, 314125, and 329930. KRP received funding from the European Research Council (ERC) under the European Union’s Horizon 2020 research and innovation programme (Grant agreement No. 866028). T. Tenkanen and G. Riddihough are acknowledged for comments on the manuscript.

## Material and Methods

### EvolveX algorithm

The flux bases of the desired traits were determined as minimum sets of fluxes requiring up- or down-regulation (respective to current metabolic phenotype) for the desired metabolic traits to arise. Such flux sets were determined by solving a mixed-integer linear programming (MILP) problem after which the required flux change directions either up or down were assigned. The MILP problem was formulated similarly to (*48*) (Equation 5).

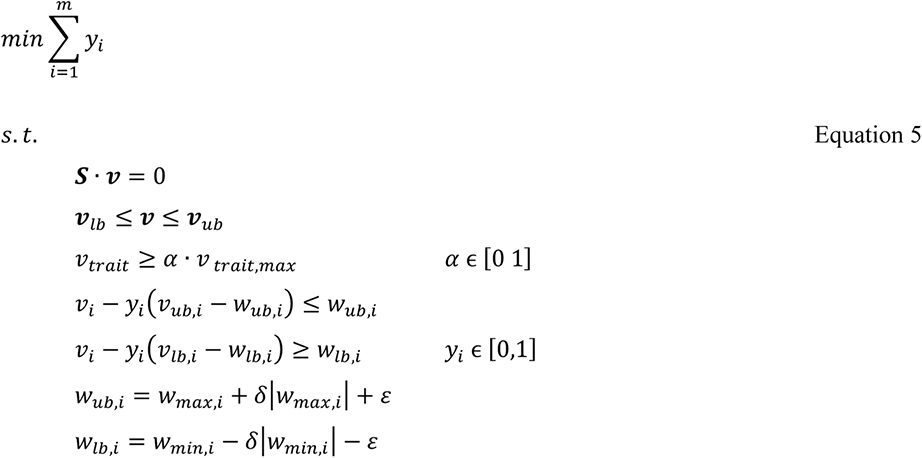

where ***S*** is the stoichiometric matrix, ***v*** the vector of metabolic fluxes, ***v***_lb_ and ***v***_ub_ the flux lower and upper bounds, respectively, *y_i_* are binary variables representing whether a reaction flux *i* requires up- or downregulation for the rise of the target trait *v_trait_* at level *α* of maximum, *v_trait,max_*. *w_ub,i_ and w_lb,_i* are the flux thresholds of wild type metabolic phenotype derived from the wild type minimum fluxes *w_min,i_* and maximum fluxes *w_max,i_* and parameters *δ* and *ε*.

As equally small, alternative, flux sets may exist, they were enumerated by introducing an additional constraint 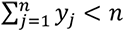 rendering previous solutions infeasible.

An evolved metabolic state was predicted by minimizing the total nutrient uptake flux for synthesizing an arbitrary unit of growth while the cells were exposed to inhibitors and metabolic effectors present in the selection niche (Equation 6).

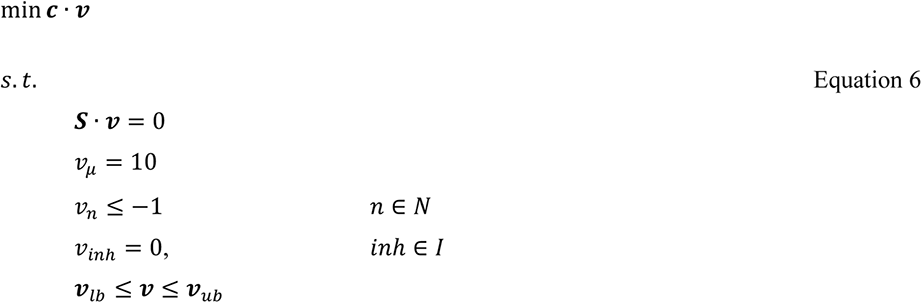

where ***c*** defines the nutrient uptakes possible in the particular selection niche with -1, *v_µ_* is the growth flux, *v_n_* are the uptake fluxes from niche *N*, and I is the set of metabolic reactions inhibited by the inhibitors present in the selection niche.

The total response to selection of the desired trait (in worst-case scenario) was predicted as the sum of growth couplings of its flux basis. For the subsets of the flux basis associated with flux change directions up and down, the minimum and maximum growth coupled fluxes, respectively, under the constraint of minimized total nutrient uptake flux for an arbitrary unit of growth, were summed (Equation 7).

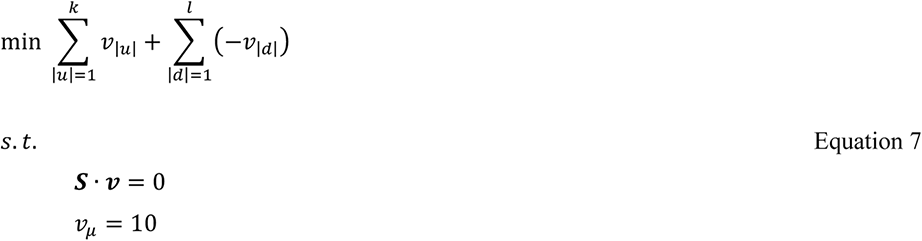

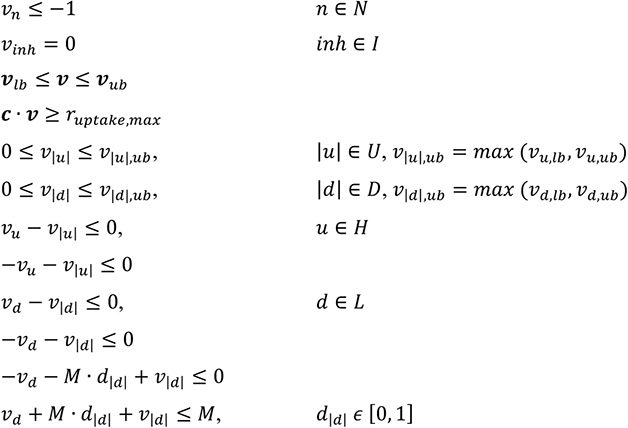

where *r_uptake,max_* is the optimal state from Equation 3, *H* and *L* are the sets of fluxes (and *k* and *l* the sizes of sets) belonging to the desired trait’s flux basis requiring up- and down-regulation, respectively, and *U* and *D* are sets of absolute flux variables representing the fluxes belonging to the desired trait’s flux basis requiring up- and down-regulation, respectively. *v_|u|,ub_* and *v_|d|,ub_* are the upper bounds of the absolute flux variables whose values were derived as maximum absolute value of the flux bounds *v_lb_* and *v_ub_*. *M* is a parameter for which a value of 20000 was used. It is double the maximum flux upper bound. *d_|d|_* is a binary variable introduced for each reversible flux belonging to the flux basis of the desired trait and requiring down-regulation.

The minimum subset size (i.e. worst-case scenario) of the flux basis of the desired trait having stronger response to selection in the particular selection niche than in common laboratory growth conditions (or target niche) was estimated under the constraint of minimized total nutrient uptake flux for an arbitrary unit of growth and under the constraint of worst-case total response to selection of the flux basis of the desired trait (Equation 8).

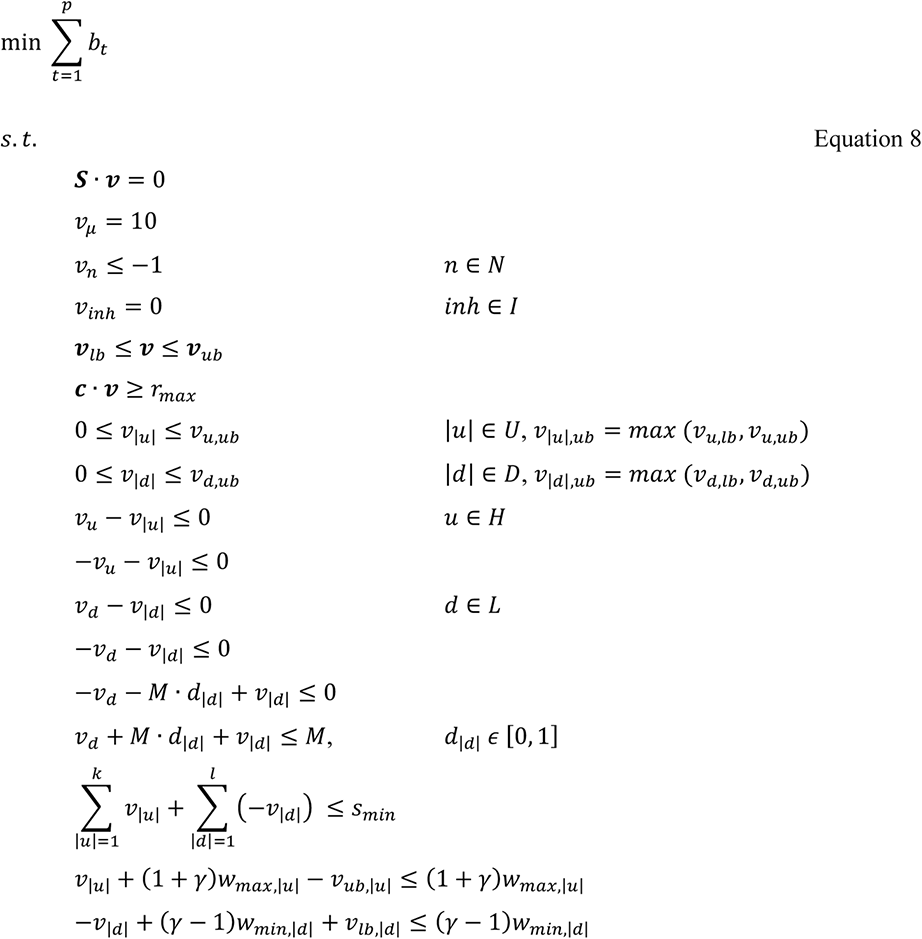

where *s_min_* is the minimum combined response to selection of the desired trait’s flux basis determined above, and *γ* is a threshold parameter for stronger response to selection than in the common laboratory conditions represented by maximum and minimum growth couplings *w_max,|u|_* and *w_min|d|_* respectively.

The suitability of an selection niche for adaptively evolving a desired metabolic trait was evaluated by deriving a weighted sum of 1) the total flux couplings to growth of the desired trait’s flux basis, 2) the minimum subset size of the desired trait’s flux basis with higher/lower predicted responses to selection than in common laboratory growth conditions (or target niche), and 3) the number of chemical components in the selection niche (Equation 9). The lower the score for the selection niche, the better suited it was considered for adaptively evolving desired traits.

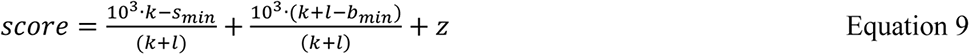

where *b_min_* is the minimum subset size (i.e. worst-case scenario) of the desired trait’s flux basis having stronger response to selection than in common laboratory growth conditions (or target niche), and *z* is the number of components in the particular selection niche.

### Model simulations

*S. cerevisiae* consensus genome-scale metabolic model v. 7.6 (*38*) was used with few revisions including augmenting the model with missing Ehrlich pathway (*42*) reactions and conditional constraints for predicting parental and evolved flux states. For implementing the model revisions, Matlab R2017b v. 9.3.0 and Cobra toolbox v.3.0 (cloned 29.03.2018) (*49*) were used. EvolveX algorithm was implemented and run in Matlab R2017b v. 9.3.0 with IBM ILOG CPLEX v. 12.8.0 functions ‘cplexlp’ and ‘cplexmilp’. The Matlab codes are available upon request.

### Strains and culture media

Both selection niches were created by modifying the defined minimal yeast growth medium by Verduyn *et al.* 1992 (*50*). The selection niche media contained 6.6 g/L K_2_SO_4_, 3 g/L KH_2_PO_4_, 0.5 g/L (MgSO_4_)7H_2_O, and the vitamins and trace elements as in Verduyn *et al.* 1992. The vitamin solution was composed of 50 mg/L d-biotin, 200 mg/L para-amino benzoic acid, 1.0 g/L nicotinic acid, 1.0 g/L Ca-pantothenate, 1.0 g/L pyridoxine-HCl, 1.0 g/L thiamine-HCl and 25 mg/L *myo*-inositol and the trace minerals solution of 3 g/L FeSO_4_·7H_2_O, 4.5 g/L ZnSO_4_·7H_2_O, 4.5 g/L CaCl_2_·6H_2_O, 0.84 g/L MnCl_2_·2H_2_O, 0.3 g/L CoCl_2_·6H_2_O, 0.3 g/LCuSO_4_·5H_2_O, 0.4 g/L NaMoO_4_·2H_2_O, 1 g/L H_3_BO_3_, 0.1 g/L KI and 15 g/LNa_2_EDTA·2H_2_O. The nitrogen and carbon sources were specific for a particular selection niche medium. The ethanol niche contained 7.5 g/L ethanol, 1.7 g/L arginine and 0.8 g/L glycine. The glycerol niche contained 5 g/L glycerol, 5g/L phenylalanine and 1.2 g/L threonine. The pH of the media were set to 6 and the media were sterile filtered.

Initially twelve different *S. cerevisiae* wine strains (commercial and vineyard isolates) were tested for their ability to grow in the two selected selection niches. The majority of the strains were able to grow sufficiently in ethanol niche after 4 days of culture, however only four strains were able to grow in glycerol niche even after a week of culture had passed. We selected as a parental strain for adaptive laboratory evolution a commercial diploid wine strain *S. cerevisiae* obtained from Lallemand which was able to grow in both selection niches.

Precultures and cells for genomic DNA extraction were grown overnight in rich medium (YPD) containing 10 g/L of yeast extract, 20 g/L peptone and 20 g/L glucose sterilized through autoclaving. Single strain isolations were performed using YPD or synthetic wine must mimicking medium (WMM) plates containing 2% agar as solidifying agent. The WMM composition was 100 g/L glucose, 100 g/L fructose, 5 g/L citric Acid, 0.5 g/L malic acid, 0.25 g/L MgSO_4_, 0.75 g/L KH_2_PO_4_, 0.5 g/L K_2_SO_4_, 0.155 g/L CaCl_2_, 0.2 g/L NaCl, 0.15 g/L NH_4_Cl, 2 mL/L anaerobic factors (1.5 g/L ergosterol, 0.5 g/L oleic acid, 50 g/L Tween 80 and 5 g/L ethanol), 3.5 mL/L amino acids solution (1.95 g/L tyrosine, 17.5 g/L tryptophan, 3.25 g/L isoleucine, 4.42 g/L aspartic Acid, 11.95 g/L glutamic Acid, 44.5 g/L arginine, 4.8 g/L leucine, 7.54 g/L threonine, 1.82 g/L glycine, 49.92 g/L glutamine, 14.56 g/L alanine, 4.42 g/L valine, 3.12 g/L methionine, 3.77 g/L phenylalanine, 7.8 g/L serine, 4.57 g/L histidine, 2.11 g/L lysine, 2.7 g/L cysteine and 59.93 g/L proline), 5 mL/L vitamins solution (2 g/L *myo*-inositol, 0.15 g/L Ca-pantothenate, 0.025 g/L thiamine-HCl, 0.2 g/L nicotinic acid, 0.036 g/L pyridoxine-HCl and 0.03 g/L biotin) and 5 mL/L trace elements solution [4 g/L MnSO_4_, 4 g/L ZnSO_4_, 1 g/L CuSO_4_, 1 g/L KI, 0.4 g/L CoCl_2_, 1 g/L H_3_BO_3_ and 1 g/L (NH_4_)Mo_7_O_24_] and the pH was set to 3.3.

### Adaptive laboratory evolution

The adaptive laboratory evolution experiment was initiated by inoculating both selection niches in triplicate from overnight single colony precultures of the parental *S. cerevisiae* strain on YPD to starting OD_600_ of 0.2. The adaptive laboratory evolution was performed for the triplicate lineages on each selection niche as a serial transfer experiment with 7 ml liquid cultures in 50 ml shake flasks at 30 °C with shaking at 180 rpm. The shake flasks were capped with cotton plugs for enhanced aeration. When culture turbidity was visually observed, the lineages were transferred to fresh medium initially to a starting OD_600_ of 0.2 and later, after the growth had improved to OD_600_ of 0.1, measured with a spectrophotometer (Ultrospec 2100, Biochrom). Intermediate lineage samples were collected to 30% w/v glycerol and stored at -80 °C.

The adaptive laboratory evolutions were initially performed for approximately 107 generations in ethanol niche and 100 generations in glycerol niche after which single colonies were picked and the isolates performing the best in the corresponding selection niche characterized as described below. The number of generations was calculated back using the following formula [Log_10_(*A_f_*/*A_i_*)]/0.3, where *A_f_* is the OD_600_ before the transfer and *A_i_* is the OD_600_ that was initially inoculated. A second round of adaptive laboratory evolution was initiated with the individual strains when they showed positive aroma profile development in the characterization (two lineages in ethanol niche, one lineage in glycerol niche), and with the last population stocks in case aroma profile improvement was not yet observed (one lineage in ethanol niche, two lineages in glycerol niche). The second round of evolution was continued approximately for additionally 97 and 65 generations in selection niches with ethanol and glycerol, respectively. Then, single colonies were again picked and the isolates performing the best in the corresponding selection niche characterized as described below.

### Characterization of evolved strains

For discarding non-genetic adaptation and for ensuring wine fermentation performance, single colonies were picked from the evolved lineages following growth on WMM + 2% agar plates for 48 h. Nine single colonies were isolated from each lineage and cultured overnight in liquid cultures on WMM. From these overnight cultures, stocks were prepared to 30% w/v Glycerol and stored at -80 °C. The overnight cultures on WMM were also used to inoculate corresponding selection niche as in adaptive laboratory evolution. Cell growth was monitored with turbidity (OD_600_) measurements. The best growing strain from each lineage was selected for a characterization in wine fermentation mimicking conditions.

Yeasts were maintained at 4 °C on YPD plates (2% glucose, 2% peptone, 1% yeast extract and 2% agar), or as glycerol stocks at −80 °C. Inocula were grown on YPD for 48 h at 25°C, washed and suspended in water.

Natural white must from the 2017 harvest was kept frozen. This must contained 215.8 g/L of sugar, density 1088.3 g/L. Enough volume for the experiment was thawed and pasteurized. Pasteurization consisted on heating to 105°C and then allowing it to cool down inside the closed autoclave.

Aliquots of 25 mL pasteurized grape must were inoculated at 0.2 final OD_600_. Fermentation was carried at 25 °C in Falcon tubes (50 ml nominal volume), capped with air locks, and monitored by weight loss. After 8 days, weight was constant, and samples were centrifuged and supernatants were kept frozen for HPLC and GCMS analysis. All experiments were performed in triplicate.

#### Determination of metabolite concentration

The concentration of glucose, fructose, glycerol, ethanol, and acetic acid was determined using a Surveyor Plus liquid chromatograph (Thermo Fisher Scientific, Waltham, MA) equipped with a refraction index and a photodiode array detector (Surveyor RI Plus and Surveyor PDA Plus, respectively) on a 300 × 7.7 mm PL Hi-Plex H+ (8 µm particle size) column (Agilent Technologies, Santa Clara, CA) and 4 x 3 mm ID Carbo-H guard (Phenomenex, Torrance, CA). The column was maintained at 50 °C and 1.5 mM H_2_SO_4_ was used as the mobile phase at a flow rate of 0.6 mL/min. Prior to injection in duplicate, the samples were filtered through 0.22 µm pore size nylon filters (Micron Analitica).

#### Analysis of volatile compounds

Samples for gas chromatography-mass spectrometry (GC-MS) analysis contained 2000 µl of sample, 1g NaCl, and 20 µl internal standard, in 20 ml flasks. Internal standard contained 1000 ppm each of 4-methyl 2-pentanol and heptanoic acid, and 100 ppm 1-nonanol, in water, prepared from 10000 ppm individual solutions in ethanol. Sample was preincubated for 10 min at 45 °C, followed by 30 min at 45 °C with 50/30 µm DBV/CAR/PDMS SPME fiber (Stableflex, SUPELCO, Bellefonte, PA). Fiber was desorbed for 5 min at 250 °C.

GC-MS was carried out in a Thermo TRACE GC Ultra apparatus coupled to a Thermo ISQ mass detector, equipped with a Thermo TriPlus autosampler. Gas chromatography was carried in a Thermo Scientific fused-silica capillary column TG-WAXMS A (30 m long; 0.25 mm OD; 0.25 µm film thickness). Chromatographic conditions were as follows: 5 min at 40 °C, 3 °C/min up to 200 °C, 15 °C/min up to 240 °C, 10 min at 240 °C. Helium was used as carrier gas at a flow rate of 1 mL/min, operating in split mode (ratio 30). Total analysis time was 71 min. Detection was performed with the mass spectrometer operating in the Full Scan mode (dwell time 500 ms), with 70 eV ionization energy, and source and quadrupole temperatures of 250°C. Detection was stopped during the time interval for ethanol elution. Peak identification was made by comparison of ion spectra with NIST mass spectral library. For each compound, including internal standards, the sum of the areas of the peaks of selected characteristic ions was obtained. Area of each compound was referred to one selected internal standard.

GC-MS data has been deposited in the Metabolights database (https://www.ebi.ac.uk/metabolights/) with accession MTBLS2208.

### Whole genome sequencing of populations and isolates

Genomic DNA was extracted from parental strains grown in YPD, lineages grown in selection niches, and single colony isolates grown in YPD using Phenol-Chloroform based extraction. Specifically, total volume of 7 mL overnight cultures was centrifuged at 3000 rpm for 3 min and the pellets were washed with sterile ddH_2_O. The cells were resuspended in 2 mL of TrisEDTA solution (0.1 M Tris and 0.1 M EDTA) and transferred to Eppendorf tubes, with 1.5 U lyticase. The pellets were then incubated at 37 °C for 30 min. Next the spheroplasted cells were centrifuged at 1500 rpm for 2 min (Eppendorf centrifuge), the supernatant was removed and the cells were resuspended in 400 µL of breaking buffer which contained 10 mM Tris, 1 mM EDTA, 100 mM NaCl, 2% Triton X-100 and 1% SDS. The cell suspensions were transferred to FastPrep Cap tubes tubes with 200 µL of glass beads (400 nm acid washed, Sigma) and the cells were broken with 3 rounds of bead beating at 4.5 Mhz/s for 20 seconds with 1 min cooling intervals or through vortexing only. The cell lysates were transferred to a new tube that contained 400 µL of phenol-chloroform/ isoamyl alchocol and 400 µL of TE buffer (Tris 50 mM, EDTA 20 mM) and were centrifuged briefly until an emulsion was formed. The emulsions were centrifuged at 13.000 rpm for 5 min at room temperature. The aqueous phase of each tube was transferred to a new Eppendorf, it was mixed with 1 mL of cold 100% ethanol and incubated at room temperature for 10 min to help precipitation. In the next step the tubes were centrifuged at 13.000 rpm for 5 min at room temperature, the ethanol was removed and the DNA pellet was resuspended at 400 µL of TE buffer with 2 µL of RNAse solution (20 mg/mL) and incubated for 15 min at 37 °C, followed by a second incubation step at 65 °C for 15 min, in order to deactivate the RNAse. The DNA solution was mixed with 400 µL of phenol-chloroform/ isoamyl alchocol and the extraction step was performed again as described above. DNA was precipitated from the aqueous phase with 1 mL of cold 100% ethanol and centrifugation at 13.000 rpm for 5 min at room temperature. The pellet was left to dry for 30 min at 55 °C, next was resuspended with 50 µL of H_2_O and was left overnight at 4 °C for the pellet to dissolve completely.

The quality of the extracted DNA was evaluated with electrophoresis in a 1% [w/v] agarose gel. DNA concentrations were measured using a Qubit (Thermo Fisher Scientific, USA). Equal amounts of DNA from all samples were used for library preparation, which was done with the NEBNext DNA Ultra2 Library Preparation Kit (New England Biolabs). The preparation of the library was performed on an automated liquid handling system (Hamilton Robotics), the quality of the library was tested on a 2100 BioAnalyzer (Agilent Technologies), and the DNA concentration was measured using a Qubit. Sequencing was performed at the Genomics Core Facility (EMBL Heidelberg) with use of the HiSeq2500 platform (Illumina, San Diego, USA) and the run produced 250 bp paired-end reads.

The sequenced samples are listed in supplementary information (Table S6) and the raw reads are deposited in ENA database (https://www.ebi.ac.uk/ena/browser/home) in study PRJEB40761 with accession numbers ERS5457098 and ERS5290477-ERS5290502 for the parental and evolved samples and ERS6110907-ERS6110952 for Panel of Normals (PoN) -samples.

### Whole genome sequence data analysis

The quality of the obtained reads was checked using Fastqc v. 0.11.4 (https://www.bioinformatics.babraham.ac.uk/projects/fastqc/; Andrews, 2010). Adapter removal and low quality read filtering was performed using cutadapt v. 1.9.1 (https://pypi.org/project/cutadapt/; Martin, 2011). The trimmed reads were aligned to *S. cerevisiae* EC1118 reference genome (*53*) with the Burrows-Wheeler Aligner v. 0.7.12 *mem* (*54*) using default parameters. The alignments were processed (added read groups, sorted, reordered, and indexed) and duplicate reads were marked using Picard Tools v. 1.129 (https://broadinstitute.github.io/picard/). Single nucleotide variant (SNV), and insertion-deletion (indel) variant calling was performed against the parental sample with GATK4 v. 4.1.0.0 (*55, 56*) *Mutect2* using the *S. cerevisiae* EC1118 as the reference and default parameters. PoN for the variant calling was compiled of 47 wild type *S. cerevisiae* strains (winery isolates and commercial wine strains, including the parental) sequenced on the same platforms as the actual samples. Variant calling was first performed for the wild type strains by running *Mutect2* in tumor-only -mode and then the panel of normal was created with GATK4 v. 4.1.0.0 *CreateSomaticPanelOfNormals*. The variant calls were filtered using GATK4 v. 4.1.0.0 *FilterMutectCalls* using default thresholds and by keeping the variants at PoN sites.

Copy number variant (CNV) analysis was performed on read counts and B-allele frequencies (baf) using GATK4 v. 4.1.0.0 tools. First, the read counts were binned for 1000 bp intervals with *CollectReadCounts*. These read counts were denoised with *DenoiseReadCounts* using the parental sample as a matched normal. The allelic counts were collected using *CollectAllelicCounts* and combined with the binned read counts for modelling the CNV segments using *ModelSegments* with number-of-changepoints-penalty-factor of eight (8). CNVs were called using *CallCopyRatioSegments*. In major copy number aberrations the default centralization of the log_2_ copy ratios to median across all contigs, misplaced the zero level to deviate from the conserved copy number. As there were major differences in the copy number aberrations between the samples, the copy ratios were re-normalized to the level of FN393086.1 contig with conserved copy number across samples. The contig was identified using the minor allele frequencies and the copy ratio differences between modelled segments. After the re-normalization, CNV’s were called for segments longer than 10 kb with -0.6 ≤ log2 copy ratio ≥ 0.3. Each CNV call was evaluated against baf data. The calls were corrected conservatively if the baf data did not support the log2 copy ratio call. The population sample calls were corrected only if the call could not be explained even by partial loss-of-heterozygosity (LoH). LoH were identified as baf zero/one segments. The log2 copy ratios of modelled segments, their copy number calls, and the LoH identified in the samples are provided in supplementary information (Table S4). Heatmap of contig median copy ratios was plotted using R v. 4.0.3 (R Core Team, 2020) *gplots* package v. 3.1.1 (*57*).

### RNA-sequencing sample preparation and data analysis

All the RNA samples were prepared according to the following procedure. Total volume of 20 mL from each culture was transferred to a 50 mL Falcon® filled with ice and was immediately centrifuged at 3000 rpm for 3 min at 4 °C (Eppendorf centrifuge). Next the supernatant was discarded and the cell pellet was snap frozen into liquid nitrogen and stored at -80 °C, until the extraction. Total RNA from the pellets was extracted with the RNAeasy kit (Qiagen) according to the manufacturer’s recommendations. In brief, 594 µL of RTL buffer plus 6 µL of β-Mercaptoethanol were used to resuspend the frozen cell pellet which was left on ice. The resuspended cells were transferred to an ice cold FastPrep Cap tube which contained 600 µL glass beads (400 nm acid washed, Sigma). The cells were then lysed with 2 cycles of bead beating, each cycle lasted 10 sec at 6 Mz/s with 15 s cooling interval. Cell lysates were transferred to a new tube and were centrifuged for 2 min at full speed (Eppendorf centrifuge) and the supernatant was carefully mixed with 1 volume of 70% HPLC-grade ethanol. Next the total volume of the sample was transferred to an RNAeasy column and the manufacturer’s instructions were followed. Total RNA was eluted with 60 µL of RNAse free water and Turbo DNAse (Invitrogen Ambion) was used to digest leftover DNA according to the manufacture instruction. Finally, one more step of RNA clean-up was performed with the same kit.

RNA library was prepared using the NEBNext® Ultra™ II Directional RNA Library Preparation Kit for Illumina: polyA transcripts capture. Briefly, barcoded stranded mRNA-seq libraries were prepared from high quality total RNA samples (∼200 ng/sample) using the NEBNext Poly(A) mRNA Magnetic Isolation Module and NEBNext Ultra II Directional RNA Library Prep Kit for Illumina (New England Biolabs (NEB), Ipswich, MA, USA) implemented on the liquid handling robot Beckman i7. Obtained libraries that passed the QC step were pooled in equimolar amounts; 2 pM solution of this pool was loaded on the Illumina sequencer NextSeq 500 and sequenced uni-directionally, generating ∼500 million reads, each 85 bases long.

The quality of the obtained RNA-sequencing reads was assessed and summarized with Fastqc v. 0.11.5 (*51*). Adapter trimming, to remove the standard lllumina TrueSeq Index adapter sequences, was performed using cutadapt v. 2.3 (*52*). Subsequently, quality read filtering and trimming was performed with FaQCs v. 2.08 (*58*), with the following parameters: *-q 20 -min_L 30 -n 3*. After trimming and filtering steps total number of reads were, in average, 31 million. Trimmed reads were then aligned to the reference genome of *S. cerevisiae* EC1118 (EnsemblFungi: annotation number GCA_000218975.1) using STAR v. 2.5.2a (*59*). On average, 85% of reads uniquely mapped to an annotated feature in the reference genome. Only uniquely mapped reads were then used to generate the gene level count tables with HTSeq v. 0.9.1(*60*). Statistical analysis was performed with R v. 3.6.1 (*61*). Differential expression analysis, including multiple testing correction and independent filtering, was performed with Bioconductor package: DESeq2 v. 1.12.0 (*62*). False discovery rate (FDR) was calculated with fdrtool v. 1.2.15 (*63*) using the raw *p*-values returned by DESeq2. Genes with a FDR < 0.05 and log2 fold change >1 or <-1 were considered as significantly differentially expressed. Unless specified, all packages were used with default parameters.

The RNA-sequencing data has been deposited to ArrayExpress database (https://www.ebi.ac.uk/arrayexpress/) with accession E-MTAB-10019.

### Proteomics sample preparation and data analysis

For the extraction of total proteome 10 mL of each culture were transferred into ice-cold 15 mL Falcon® tubes which were centrifuged immediately at 3000 rpm for 3 min at 4 °C (Eppendorf centrifuge). The supernatant from the centrifugation was discarded and the cell pellets were washed once with 1 mL of cold PBS buffer. The washed pellets were snapped frozen with liquid nitrogen and stored at -80 °C. For the extraction, the cell pellets were lysed with 0.1% RapiGest (Waters) in 100 mM ammonium bicarbonate, followed by mechanical disruption with 3 rounds of sonication (1 cycle: 10 sec sonication and 10 sec rest on ice per round). Sonication was followed by 2 cycles of bead beating (200 µL glass beads, 400 nm acid washed, Sigma), each cycle lasting 20 sec at 4 Mz/sec with 1 min cooling intervals between the cycles.

Reduction of disulphide bridges in cysteine containing proteins was performed with dithiothreitol (56 °C, 30 min, 10 mM in 50 mM HEPES, pH 8.5). Reduced cysteines were alkylated with 2-chloroacetamide (room temperature, in the dark, 30 min, 20 mM in 50 mM HEPES, pH 8.5). Samples were prepared using the SP3 protocol (*64*) and trypsin (sequencing grade, Promega) was added in an enzyme to protein ratio 1:50 for overnight digestion at 37 °C. Peptides were labelled TMT10plex (*65*) Isobaric Label Reagent (ThermoFisher) according the manufacturer’s instructions. For further sample clean up an OASIS® HLB µElution Plate (Waters) was used. Offline high pH reverse phase fractionation was carried out on an Agilent 1200 Infinity high-performance liquid chromatography system, equipped with a Gemini C18 column (3 µm, 110 Å, 100 x 1.0 mm, Phenomenex), (*66*), resulting in 12 fractions.

After fragmentation, the peptides were separated using an UltiMate 3000 RSLC nano LC system (Dionex) fitted with a trapping cartridge (µ-Precolumn C18 PepMap 100, 5µm, 300 µm i.d. x 5 mm, 100 Å) and an analytical column (nanoEase™ M/Z HSS T3 column 75 µm x 250 mm C18, 1.8 µm, 100 Å, Waters). Trapping was carried out with a constant flow of trapping solution (0.05% trifluoroacetic acid in water) at 30 µL/min onto the trapping column for 6 min. Subsequently, peptides were eluted via the analytical column running solvent A (0.1% [v/v] formic acid in water) with a constant flow of 0.3 µL/min, with increasing percentage of solvent B (0.1% [v/v] formic acid in acetonitrile) from 2% to 4% in 4 min, from 4% to 8% in 2 min, then 8% to 28% for a further 37 min, in another 9 min. from 28%-40%, and finally 40%-80% for 3 min followed by re-equilibration back to 2% B in 5 min. The outlet of the analytical column was coupled directly to an Orbitrap QExactive™ plus Mass Spectrometer (Thermo) using the Nanospray Flex™ ion source in positive ion mode.

The peptides were introduced into the QExactive plus via a Pico-Tip Emitter 360 µm OD x 20 µm ID; 10 µm tip (New Objective) and an applied spray voltage of 2.2 kV. The capillary temperature was set at 275 °C. Full mass scan was acquired with mass range 375-1200 m/z in profile mode with resolution of 70000. The filling time was set at maximum of 100 ms with a limitation of 3x10_6_ ions. Data dependent acquisition (DDA) was performed with the resolution of the Orbitrap set to 17500, with a fill time of 50 ms and a limitation of 2x10_5_ ions. A normalized collision energy of 32 was applied. Dynamic exclusion time of 20 s was used. The peptide match algorithm was set to ‘preferred’ and charge exclusion ‘unassigned’, charge states 1, 5 - 8 were excluded. MS data was acquired in profile mode.

The acquired data were processed using IsobarQuant (*67*) and Mascot v. 2.2.07. A Uniprot *Saccharomyces cerevisiae* proteome database (UP000002311) containing common contaminants and reversed sequences was used. The search parameters were the following: Carbamidomethyl (C) and TMT10 (K) (fixed modification), Acetyl (N-term), Oxidation (M) and TMT10 (N-term) (variable modifications). A mass error tolerance of 10 ppm was set for the full scan (MS1) and for MS/MS (MS2) spectra of 0.02 Da. Trypsin was selected as protease with an allowance of maximum two missed cleavages. A minimum peptide length of seven amino acids and at least two unique peptides were required for a protein identification. The false discovery rate on peptide and protein level was set to 0.01. GO-process enrichments were determined using https://www.yeastgenome.org/goTermFinder with the metabolic enzymes annotated in *S. cerevisiae* consensus genome-scale metabolic model v. 7.6 (*38, 68*) at *p*-value 0.1.

The mass spectrometry proteomics data have been deposited to the ProteomeXchange Consortium via the PRIDE (*69*) partner repository with the dataset identifier PXD023171.

### Small scale fermentation of natural wine must (microvinification) for transcriptomics and proteomics analysis

A single colony of the parental strain, two evolved isolates originating from the ethanol niche and two evolved isolates originating from the glycerol niche were grown overnight in 50 mL Falcon^®^ tubes with 15 mL of YPD. The overnight grown cells were washed three times with PBS and diluted to an initial OD_600_ of 0.1 in 55 mL of natural white must from the 2017 harvest (see above). For the microvinification process, 50 mL Erlenmeyer flaks were used, filled to the maximum, in order to create microanaerobic conditions. Maintaining the anaerobic conditions meant that the growth could not be estimated based on changes in the optical density, but it was correlated with the observed weight loss, which occurs from the release of CO_2_, the end product of carbon metabolism. Release of CO_2_ is possible through a small needle which is pierced through rubber plugs, which in turn were sterilized and used to seal the Erlenmeyer flaks, while a small piece of gauge prevents anything from the environment to fall inside the flask through the needle. The growth stage of the cultures was estimated based on weight loss which correlates to the consumption of glucose and release of CO_2_ as suggested by Harsch *et al.* (2010). For this reason, the initial weight of the cultures was measured and followed once every day until no more weight loss was observed, at which point the cultures had entered stationary phase. After the establishment of the growth kinetics with weight loss, same cultures as described above were prepared, weight loss was once again followed and the cells were harvested at mid exponential phase for RNA-sequencing and proteomics analysis, as described at the previous sections.

### Data analysis and visualization

Principal component analysis was performed using R v. 4.0.3 (*71*) and *factoextra* package v. 1.0.7 (*72*). Data processing was performed using *readr* package v. 1.4.0 (*73*), *dplyr* package v. 1.0.2 (*74*) and *tidyr* package v. 1.1.2 (*75*). For statistical plotting *ggplot2* package v. 3.3.2 was used, the euler diagram was generated using *eulerr* package v. 6.1.0 (*76*), CNV heatmap was plotted using heatmap.2 function from *gplots* package v. 3.1.1 (*57*), and the color schemes were obtained from RColorBrewer package v. 1.1-2 (*77*), and *ggsci* package v. 2.9 (*78*).

## Supplementary information

Table S1: EvolveX scores enumerated for selection niche suitability for adaptively evolving the target aromas

Table S2: Flux bases and surrogate traits

Table S3. Performance in wine must fermentations

Table S4. Aneuploidy and LoH

Table S5. Significantly differentially abundant metabolic enzymes in selection niche and in wine must fermentation and corresponding model predictions

Table S6. Sample identifiers

